# Flavinated SDHA Underlies the Change in Intrinsic Optical Properties of Oral Cancers

**DOI:** 10.1101/2023.07.30.551184

**Authors:** Tomoko Marumo, Chima V. Maduka, Evran Ural, Ehsanul Hoque Apu, Seock-Jin Chung, Nynke S. van den Berg, Quan Zhou, Brock A. Martin, Eben L. Rosenthal, Takahiko Shibahara, Christopher H. Contag

**Affiliations:** Department of Oral and Maxillofacial Surgery, Tokyo Dental College, 2-9-18 Kanda-Misakicho, Chiyoda-ku, Tokyo 101-0061, Japan; Department of Biomedical Engineering, Michigan State University, East Lansing, MI 48824, USA; Institute for Quantitative Health Science & Engineering, Michigan State University, East Lansing, MI 48824, USA; Comparative Medicine & Integrative Biology, Michigan State University, East Lansing, MI 48824, USA; Division of Hematology and Oncology, Department of Internal Medicine, Michigan Medicine, University of Michigan, Ann Arbor, MI 48109, USA; Department of Otolaryngology – Division of Head and Neck Surgery, Stanford University School of Medicine, 269 Campus Drive, Stanford, CA 94305, USA; Department of Pathology, Stanford University School of Medicine, 3100 Pasteur Drive, Stanford, CA 94305, USA; Department of Otolaryngology – Head and Neck Surgery, Vanderbilt University Medical Center, 1211 Medical Center Dr, Nashville, TN 37232; Department of Microbiology & Molecular Genetics, Michigan State University, East Lansing, MI 48824, USA

**Keywords:** Autofluorescence, oral squamous cell carcinoma (OSCC), flavin adenine dinucleotide (FAD), succinate dehydrogenase, metabolism.

## Abstract

The molecular basis of reduced autofluorescence in oral squamous cell carcinoma (OSCC) cells relative to normal cells has been speculated to be due to lower levels of free flavin adenine dinucleotide (FAD). This speculation, along with differences in the intrinsic optical properties of extracellular collagen, lie at the foundation of the design of currently-used clinical optical detection devices. Here, we report that free FAD levels may not account for differences in autofluorescence of OSCC cells, but that the differences relate to FAD as a co-factor for flavination. Autofluorescence from a 70 kDa flavoprotein, succinate dehydrogenase A (SDHA), was found to be responsible for changes in optical properties within the FAD spectral region with lower levels of flavinated SDHA in OSCC cells. Since flavinated SDHA is required for functional complexation with succinate dehydrogenase B (SDHB), decreased SDHB levels were observed in human OSCC tissue relative to normal tissues. Accordingly, the metabolism of OSCC cells was found to be significantly altered relative to normal cells, revealing vulnerabilities for both diagnosis and targeted therapy. Optimizing non-invasive tools based on optical and metabolic signatures of cancers will enable more precise and early diagnosis leading to improved outcomes in patients.

---

Early detection of oral squamous cell carcinoma (OSCC) is characterized by a 5-year survival rate exceeding 80%, whereas late diagnosis leads to less than 38% survival rates^1–3^, underlying the vital relationship between prompt diagnosis and outcome in patients. Currently, clinical diagnostic methods include palpation and visual inspection with, or without, the aid of biological reagents, such as toluidine blue or Lugol’s iodine^4, 5^. While being helpful, chemical agents could cause allergies and visual inspection is incapable of delineating pre-neoplastic tissues^6^, informing the need for label-free noninvasive diagnostic tools, including optical detection of tissue autofluorescence.

Tissue autofluorescence for early detection of OSCC is based on the observation that cancers exhibit lowered levels of endogenous fluorescent molecules; these could include flavin adenine dinucleotide (FAD), crosslinked collagen and reduced nicotinamide adenine dinucleotide (NADH)^7–9^. Consequently, autofluorescent intensity is reduced in dysplastic and neoplastic tissues in comparison to surrounding normal tissues when exposed to light in the ultraviolet (UV) and blue regions of the spectrum^10^. Exploiting this feature, handheld devices have been developed for visualizing the spectral properties of FAD or collagen, and several of these have been commercialized for screening for OSCC in patients^11^. Evaluating the loss of tissue autofluorescence in OSCC lesions is currently subjective due to lacking uniform standards; the biochemical basis of signal loss remains, in part, inferred from the spectral properties of free FAD. Furthermore, although FAD is an important metabolic cofactor^9^, the functional metabolic profile of OSCC is yet to be fully elucidated and related to autofluorescence.

Here, we elucidate the molecular basis of decreased tissue autofluorescence in OSCC cells which is presumed to arise from altered levels of FAD, and characterize underlying functional metabolism. To exclude the contribution of collagen, angiogenesis and accompanying hemoglobin, as well as heterogenous morphological changes in autofluorescence^12–14^, we first analyzed human-derived cultured cells. We show that commercial optical instruments which rely on the spectral properties of FAD may not be optimal for detecting OSCC. Instead, we identify unique spectral signatures that could be targeted to increase signal-to-noise ratios. Second, we confirm varying levels of loss in autofluorescence among OSCC cell lines compared to non-cancer cells, and show that differential autofluorescence could not be accounted for by free FAD alone. By integrating spectral and proteomic analyses, we identify succinate dehydrogenase subunit A (SDHA), a 70 kDa protein, in the FAD spectral region to account for altered optical properties of OSCC. Since flavinated SDHA is required for functional complexation with succinate dehydrogenase B (SDHB), we evaluated levels of SDHB in tissues from cancer patients as a functional measure of levels of flavinated SDHA, and observed decreased SDHB expression in human OSCC tissues compared to normal regions of the patient-derived tissue samples. Third, we observed a unique metabolic phenotype, characterized by increases in glycolytic flux and oxidative phosphorylation, underlying loss of autofluorescence in OSCC. Inhibition of oxidative phosphorylation is not accompanied by bioenergetic compensation, revealing a therapeutic opportunity that targets the metabolic vulnerabilities of OSCC. Optimizing non-invasive tools used for detecting OSCC will enable earlier and more precise diagnosis than is currently possible, which will positively correlate with good prognosis in OSCC patients.

## Results

### Cancer detection based on spectral properties of free FAD is not optimal for oral squamous cell carcinoma (OSCC)

We used a commonly-applied, handheld, screening device for oral cancer (excitation: 400-460 nm, emission: 470-580 nm)^15^. The design of this device is based on the use of safe ultraviolet and blue light illumination, and the spectral properties of free FAD^16^. It identifies dark areas due to reduced autofluorescence of neoplastic tissues; however, other areas which may, or may not, correspond to neoplasia are also identified (Fig. 1a-b). To validate that cell lines can model changes in tissue autofluorescence, non-cancer, epithelial (squamous) HaCaT cell lines were selected as controls for oral squamous cell carcinoma cell lines. We assessed the spectral characteristics of non-cancer (HaCaT) and human-derived OSCC (Ca9-22, HSC-3 and SAS) cell lysates over a range of wavelengths that included those applied by optical devices. This range of wavelengths are denoted by red dashed rectangles on the fluorescence excitation-emission matrices (EEMs) of cell lysates, along with purified FAD and NADH (Fig. 2a-j). For reference, the EEMs of FAD and NADH were measured and confirm that this optical device measures FAD (Fig. 2a) and not NADH (Fig. 2b) autofluorescence. In addition, the EEM of FAD and NADH reveal their bimodal and unimodal spectra, respectively (Fig. 2a-b). There was less autofluorescence among OSCC relative to the HaCaT cells (Fig. 2c-f), consistent with observations in tissues (Fig. 1a-b). Subtracting the EEM of OSCC lysates from the EEM of HaCaT cell lysates, and considering first and second order scattering, reveal the possible sources of autofluorescence (Fig. 2g-j), and suggests that the selected wavelengths are not optimal. The use of cell lysates was intended to reduce the contribution of cellular and subcellular structures to the scatter. We identified spectral regions at excitation (360-400 nm) and emission (575-650 nm) where optical signals from OSCC lysates dramatically differ from that of non-cancer cells (Fig. 2i-j).

**Figure 1.**
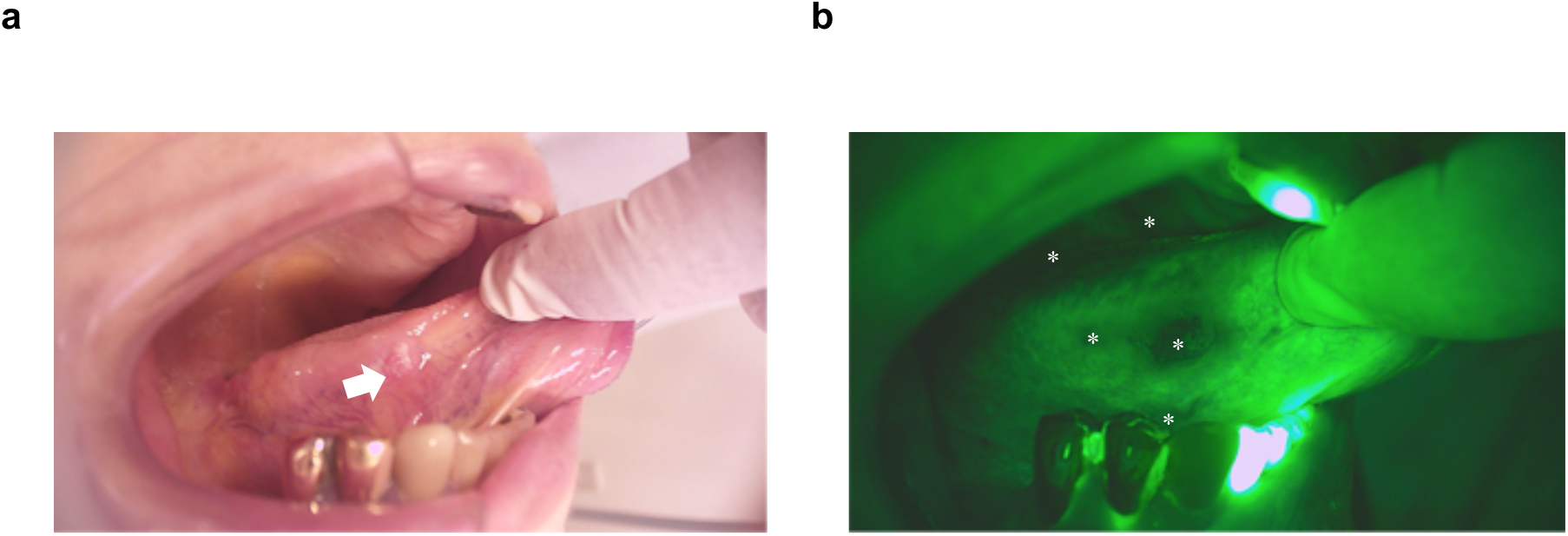
Commercial handheld devices for detection of oral squamous cell carcinoma (OSCC) are based on loss of flavin adenine dinucleotide (FAD) autofluorescence, but it is not clear if this is free or bound FAD and whether commercial devices have been optimized for the molecular basis of loss of fluorescence. **a**, Malignant lesion (white arrow) on the tongue of a patient having OSCC. **b**, Visualization of neoplastic lesions (white asterisks) using a handheld device reveals a loss of fluorescence in multiple areas (white letters), which may or may not correspond to definitive lesions; the handheld device is based on loss of free FAD autofluorescence.

**Figure 2.**
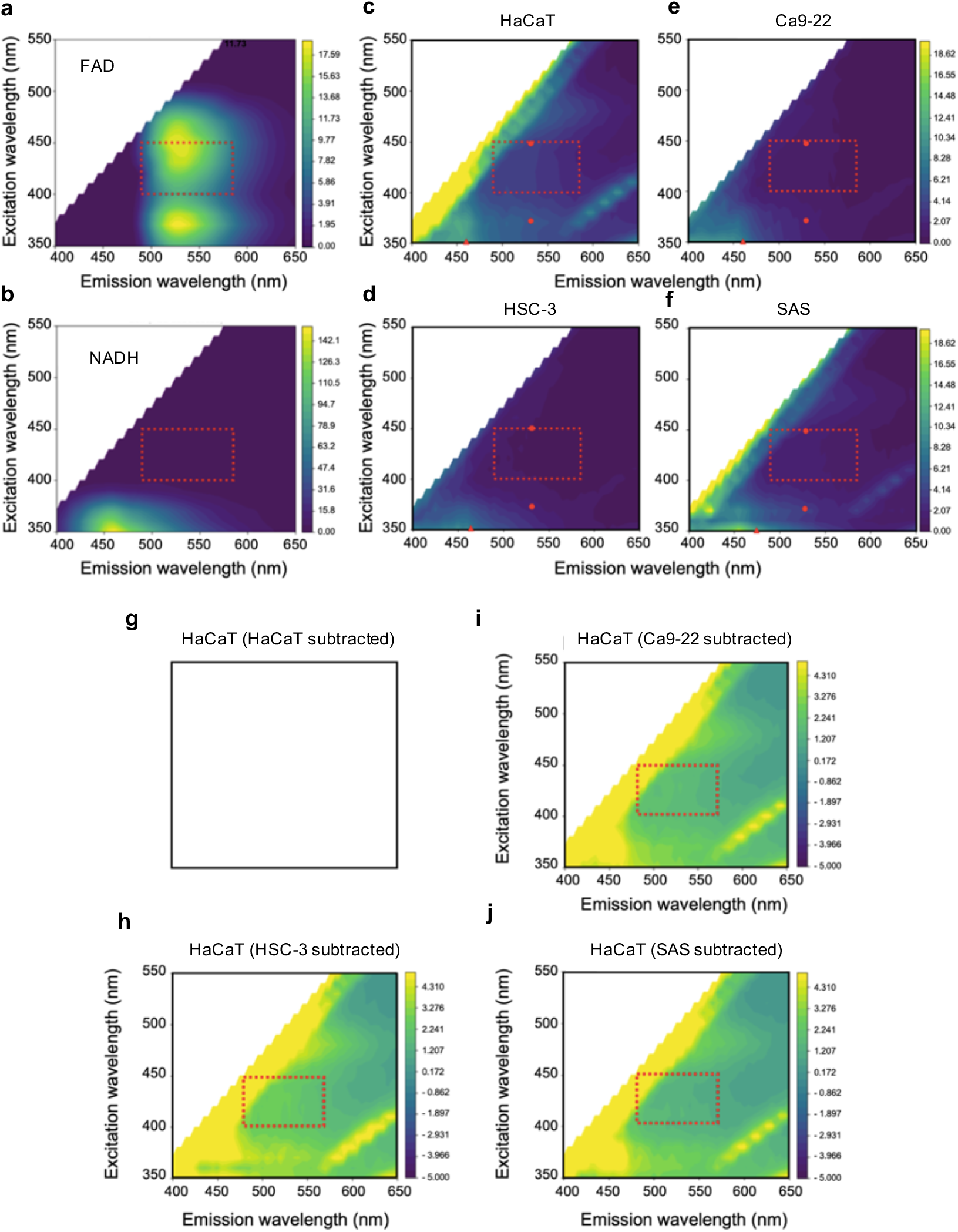
Fluorescence excitation-emission matrices (EEMs) suggest a loss of intensity in oral squamous cell carcinoma (OSCC) cell lysates but also demonstrate that the EEMs of free flavin adenine dinucleotide (FAD) are not optimal for OSCC detection. **a**, The fluorescence EEM of free FAD is characteristically bimodal, having two emission peaks at 530 nm with different (360 nm, 445 nm) excitation wavelengths. **b**, The EEM of reduced nicotinamide adenine dinucleotide (NADH) is unimodal (excitation at 360 nm, emission at 460 nm); the red dashed rectangles indicate the range of wavelengths (of free FAD) observable with handheld devices, and each contour corresponds to a region of equal fluorescence intensity. **c-f**, In comparison to non-cancer (HaCaT; **c**) lysates, representative EEM spectra (quantified in Fig. 3a) suggest loss of intensity in OSCC cell lysates (**d-f**) but there are more non-fluorescent than fluorescent regions within the red dashed rectangles; red dots and triangles correspond to the peaks for FAD and NADH, respectively. **g,** Simulated representation of subtracting EEMs of HaCaT lysates from itself. **h-j**, Subtracting the EEMs of OSCC from the EEM of HaCaT cell lysates reveals that there are regions in the EEMs that are not within red dashed rectangles; additionally, there is a spectral region at excitation (360-400 nm) and emission (575-650 nm) that is distinct in all OSCC lysates.

### Flavinated succinate dehydrogenase subunit A (SDHA) is responsible for differential autofluorescence in OSCC cells

Size-exclusion ultrafiltration experiments were undertaken comparing HaCaT to SAS cells, where autofluorescence was measured in cell suspension, unfiltered supernatant obtained after cell lysis, and filtered supernatant using 30 or 10 kDa ultra filters. Interestingly, a decrease in autofluorescence was observed in SAS compared to HaCaT in the cell suspension, unfiltered supernatant and the supernatant filtered using 30kDa ultra filters (Supplementary Fig. 1a-c). However, the differential change in autofluorescence was lost after ultrafiltration with the 10 kDa ultra filter (Supplementary Fig. 1d); these data suggested that the molecule causing the autofluorescence is greater than 10 kDa in size, excluding small molecules, such as FAD and NADH. Therefore, we analyzed cell lysates using a modified form of sodium dodecylsulfate polyacrylamide gel electrophoresis (SDS-PAGE) to identify the source of cellular autofluorescence. Blue light excitation of unstained SDS-PAGE gels revealed one distinct autofluorescent band around 70 kDa in HaCaT lysates as being responsible for altered levels of autofluorescence (Fig. 3a). Quantitating fluorescence on the SDS-PAGE gels showed a 2.3-, 3- and 1.8-fold decrease in OSCC (Ca9-22, HSC-3 and SAS, respectively) compared to HaCaT lysates (Fig. 3a). To further characterize the autofluorescent protein(s) in the SDS-PAGE gel, we performed liquid chromatography with tandem mass spectrometry (LC-MS/MS) on the band that migrated to approximately 70 kDa, and determined that there were 16 candidate proteins present in HaCaT that were present at lower levels in OSCC cell lysates (Supplementary Table 1). High-resolution autofluorescence imaging of HaCaT and OSCC cells revealed cellular localization and a characteristic punctate autofluorescent pattern in the cytoplasm, allowing us to narrow down our analyses to protein candidates that are enriched in organelles^17^ (Supplementary Fig. 2a-b). Flavinated proteins are a source of autofluorescence and commonly present in mitochondria^18, 19^. Two proteins that met the criteria of being approximately 70 kDa mitochondrial flavoproteins were very long-chain specific acyl-CoA dehydrogenase (ACADV) and SDHA (Supplementary Table 2). Accordingly, we probed the autofluorescent protein(s) in the SDS-PAGE gel by western blot for SDHA. We observed that SDHA levels were reduced by 2.6- and 2.1-fold in Ca-922 and SAS relative to HaCaT cell lysates (Fig. 3b). In contrast, although ACADV was detected (Fig. 3c), its levels were variable, increasing in some OSCC cell lysates (Fig. 3c), with GADPH levels being similar between groups (Fig. 3d). Imaging by phase contrast and immunofluorescence reinforced cytoplasmic localization of both ACADV and SDHA (Fig. 4a).

**Figure 3.**
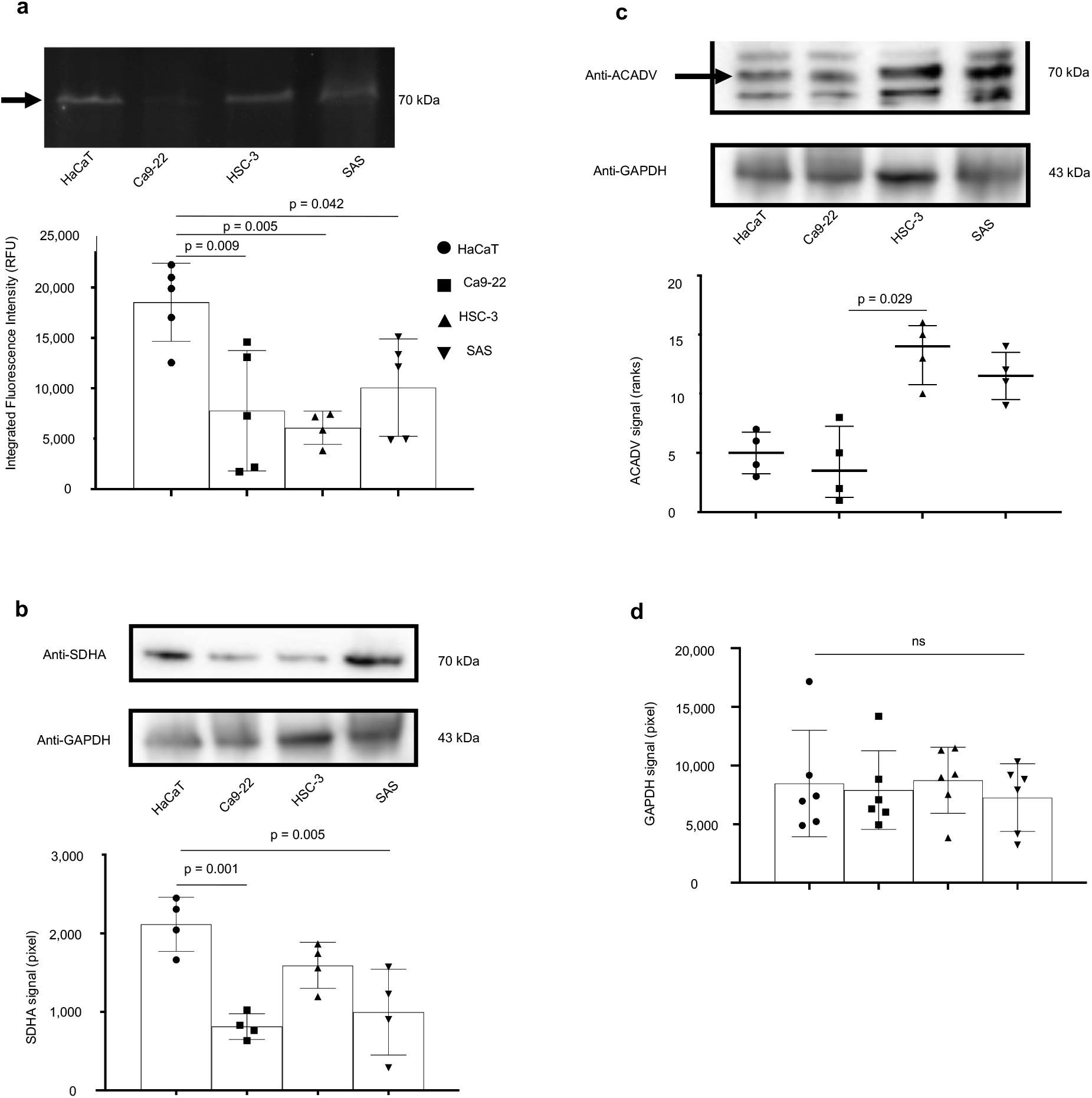
Flavinated succinate dehydrogenase complex subunit A (SDHA) accounts for lowered fluorescence intensity in oral squamous cell carcinoma (OSCC) compared to non-cancer cells. **a**, Representative SDS-PAGE gel and quantitation when exposed to blue light suggest that decreased autofluorescence occurs at the 70 kDa region, and that fluorescence intensity is lower in OSCC cell (SAS, HSC-3 and Ca9-22) lysates compared to non-cancer (HaCaT) lysates. **b**, Representative western blot image of proteins transferred from SDS-PAGE gel shows SDHA is detected in the 70 kDa region where autofluorescence was previously observed; also, SDHA levels decrease in OSCC compared to HaCaT cells. **c**, Western blot image of proteins transferred from SDS-PAGE gel shows ACADV is also detected at the 70 kDa region where autofluorescence was previously observed; however, ACADV levels do not decrease in OSCC compared to HaCaT cells. **d**, GAPDH levels in western blots are similar between OSCC and HaCaT cells. Not significant (ns); Kruskal-Wallis test followed by Dunn’s multiple comparison test was used to assess ranks expressed as median (interquartile range) in Fig. 3f; Other datasets were analyzed by one-way ANOVA followed by Tukey’s multiple comparison test and expressed as mean (SD); Grubb’s test eliminated an outlier in the HSC-3 group for Fig. 3a; n = 4-5 (Fig. 3a), n = 4 (Fig. 3d, f), n = 6 (Fig. 3g).

**Figure 4.**
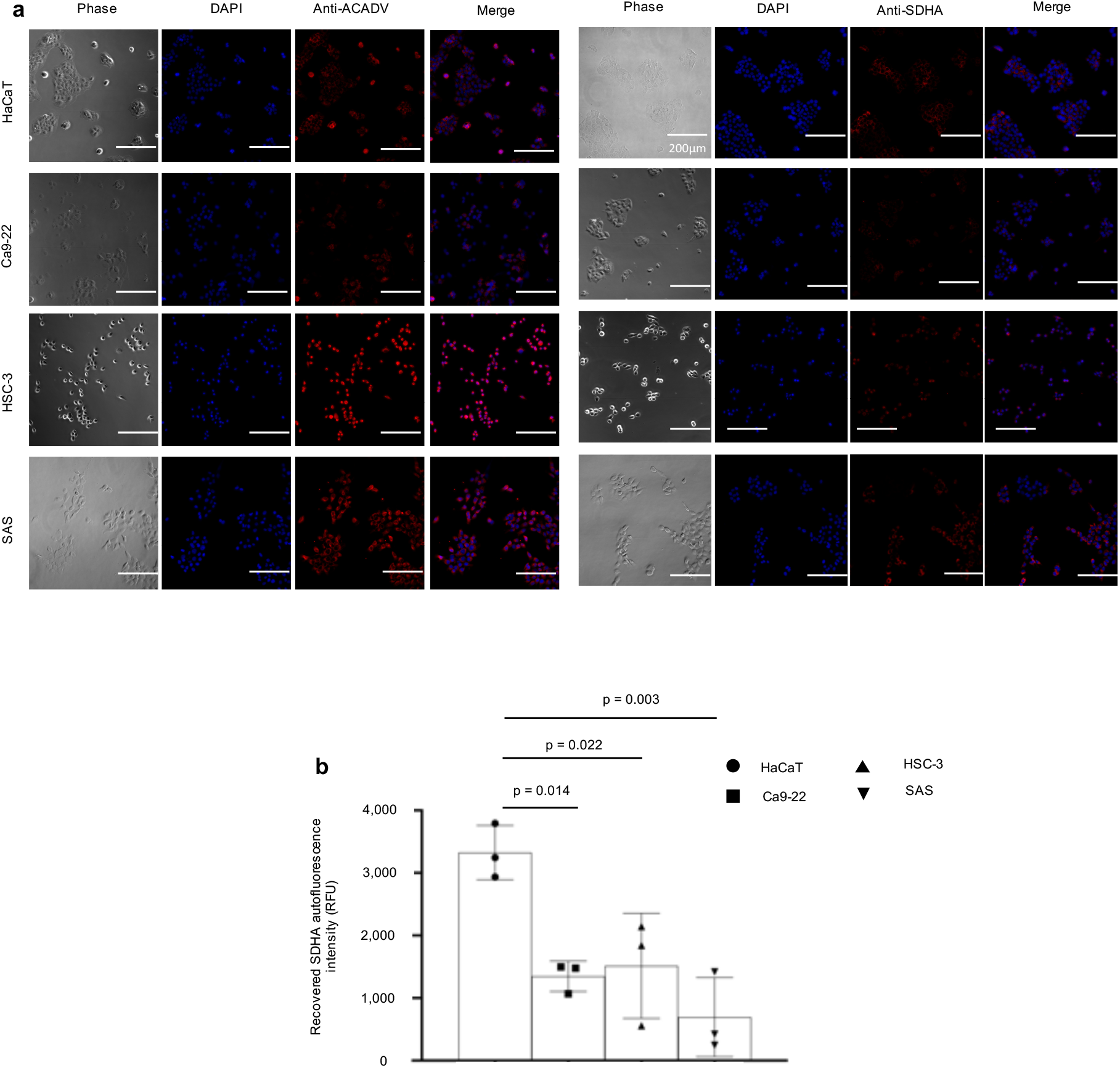
SDHA expression is reduced in oral squamous cell carcinoma (OSCC) cell lines, and both SDHA and ACADV are localized to the cytoplas. **a**, Representative phase contrast and immunofluorescent images reveal cytoplasmic localization of ACADV and SDHA in cell lines; scale bars are 200 μm. **b**, Relative fluorescence intensity of SDHA complexes recovered from cell lysates via immunoprecipitation with magnetic beads reveals decreased SDHA expression in OSCC (Ca9-22, SAS, HSC-3) when compared to non-cancer (HaCaT) cells.. One-way ANOVA followed by Tukey’s multiple comparison test, mean (SD), n = 3.

To substantiate the relationship between autofluorescence levels and SDHA in these human cell lines, we used magnetic bead affinity purification to isolate intracellular SDHA from cell lysates and measured the amounts of autofluorescent (i.e. flavinated) SDHA that could be recovered from these cell lines. Under blue light, the fluorescence from SDHA isolated by immunoprecipitation corroborated decreased autofluorecent signal intensities for Ca9-22, HSC-3 and SAS by 2.5-, 2.2- and 4.8-fold, respectively, compared to HaCaT cells (Fig. 4b). The apparent lower levels of flavinated SDHA in OSCC cell lines suggested a need to assess flavinated SDHA levels in human OSCC tissues relative to surrounding normal tissues to determine the translational significance of our observations. Cancer and normal regions in tissue sections from oral cancer patients were delineated by a board-certified pathologist, and comparisons of SDHA and succinate dehydrogenase subunit B (SDHB) levels in each region were determined (Fig. 5a). SDHB forms complexes with only flavinated SDHA for mammalian, functional, enzymatic activity^20^. Consequently, SDHB levels could be a functional readout for flavinated SDHA, reflecting levels of the flavo protein^20^. As anticipated, overall SDHA levels were similar among the cell types (Fig. 5a; quantitative data not shown), but SDHB levels were 1.8-fold lower in cancer compared to adjacent normal tissues (Fig. 5b).

**Figure 5.**
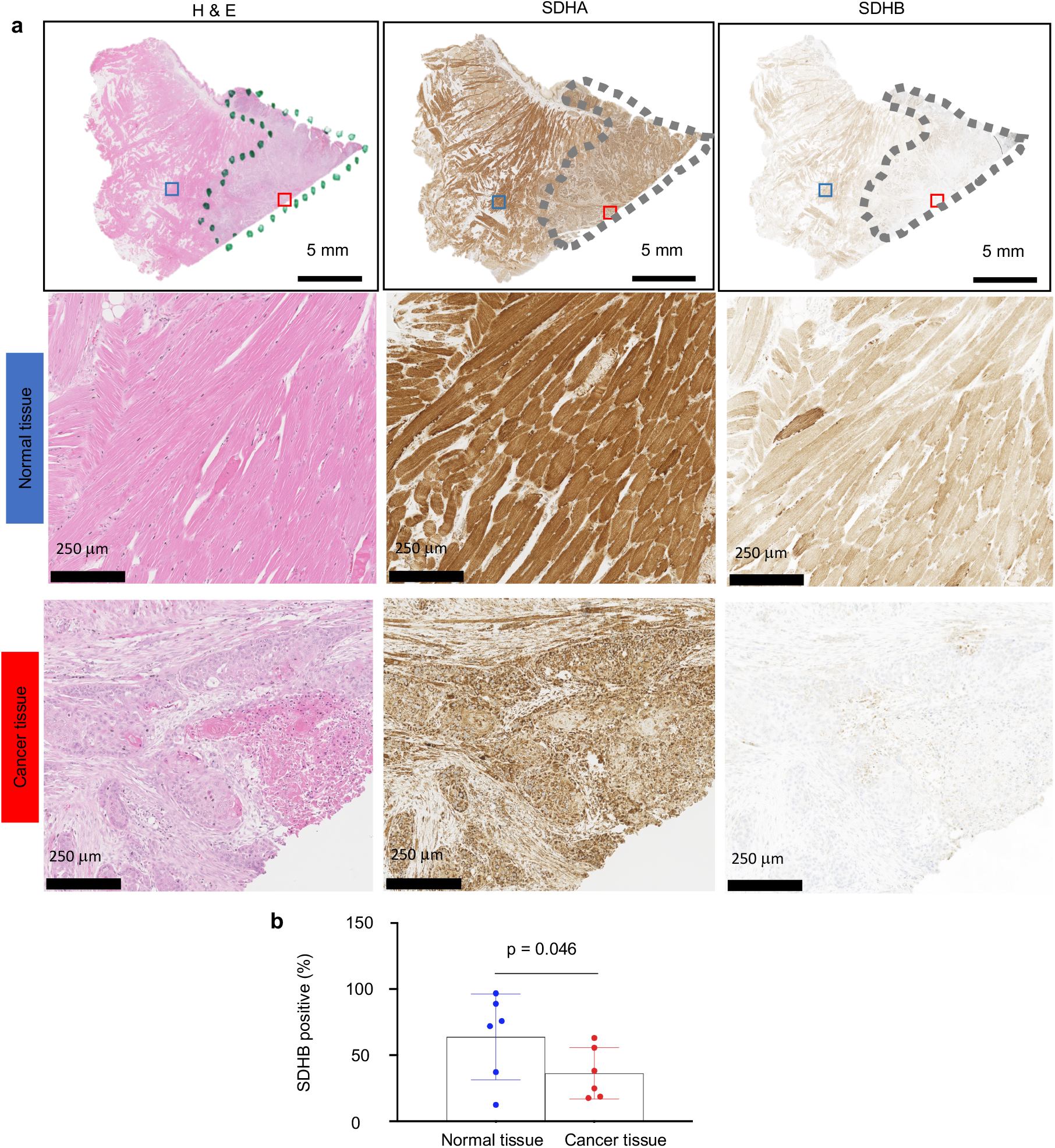
Succinate dehydrogenase complex subunit B (SDHB) levels are reduced in oral squamous cell carcinoma (OSCC) tissues. **a**, Representative hematoxylin and eosin (H & E), SDHA and SDHB immunohistochemical staining of cancer (outlined by dotted lines) and normal tissues; higher magnification of normal and cancer tissues in blue and red squares, respectively, suggest loss of both SDHA and SDHB expression. **b**, Quantified levels of SDHB are lower in cancer compared to normal tissues. Two-tailed paired t-test, n = 6 (refer to methods section).

### Altered metabolism underlies loss of autofluorescence in OSCC

To assess the metabolic underpinnings of changing autofluorescence in OSCC, likely reflected in the altered levels of flavinated SDHA, we measured oxygen consumption rate (OCR), extracellular acidification rate (ECAR) and lactate-linked proton efflux rate (PER), indices of oxidative phosphorylation, glycolytic flux and monocarboxylate transporter function^21^, respectively, used in evaluating metabolic reprogramming^22, 23^. Compared to HaCaT cells, OCR is elevated in HSC-3 and SAS by 1.8- and 2.8-fold, respectively (Fig. 6a); ECAR is increased in HSC-3 and SAS lines by 2.4- and 2.1-fold, respectively (Fig. 6b); and PER is increased in HSC-3 and SAS lines by 2.4- and 2.1-fold, respectively (Fig. 6c). Furthermore, inhibition of complex V (ATP synthase) of the mitochondrial electron transport chain does not completely account for differential oxygen consumption; additional inhibition of complex I and III is required (Fig. 6d). Of note, there is no compensatory elevation in ECAR or PER following inhibition of oxygen consumption by rotenone and antimycin A (Fig. 6e-f).

**Figure 6.**
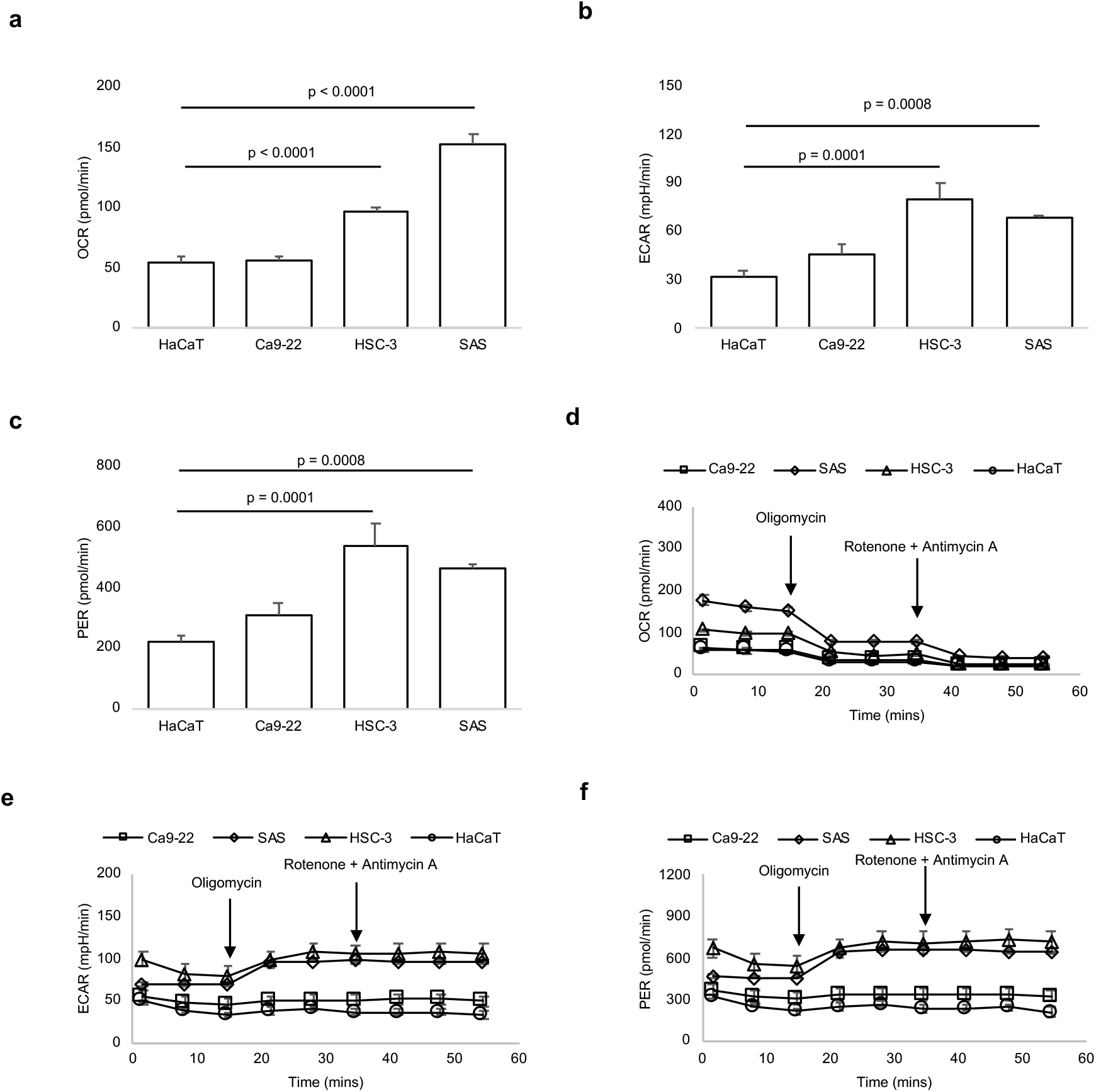
Altered metabolism underlies loss of tissue autofluorescence. **a-c**, Compared to non-cancer (HaCaT) cells, basal oxygen consumption rate (OCR; **a**), extracellular acidification rate (ECAR; **b**) and proton efflux rate (PER; **c**) are elevated in HSC-3 and SAS cancer cell lines. **d**, Differential OCR levels in SAS and HSC-3 are not accounted for by only inhibiting ATP synthesis (synthase) using oligomycin. **e-f**, There is no compensatory increase in ECAR (**e**) or PER (**f**) in SAS and HSC-3 after inhibiting complex I and III using rotenone and antimycin A, despite reduction in OCR. One-way ANOVA followed by Tukey’s multiple comparison test, mean (SD), n = 3 biological and n = 3 technical replicates.

## Discussion

Oral cancer screening is critical to the early detection of these malignancies. As noted by the American Dental Association Council on Scientific Affairs, tools for effective early detection of oral cancer remains an unaddressed need^24^. Commercial handheld optical devices for oral squamous cell carcinoma (OSCC) screening are often based on lowered levels of free flavin adenine dinucleotide (FAD) in cancer cells. Lowered FAD levels manifest as decreased autofluorescence (excitation peaks: 360 nm or 445 nm, emission peak: 530 nm) relative to what is observed in surrounding normal tissues. Our findings illustrate that such optical devices may not be optimized for the detection of OSCC. Such optical devices could be improved by tuning the excitation source and emission windows to 430-450 nm and 480-500 nm, respectively, since OSCC cell lysates have reduced signals in these regions of the spectral profiles. However, contributions of scattering in these regions of the spectrum and the small Stokes shifts may still result in poor signal-to-noise ratios and suboptimal device performance^25^. To identify optimal parameters, we subtracted the excitation-emission matrices (EEM) of OSCC cell lysates from non-cancer cell lysates to delineate distinct autofluorescence and scatter profiles. This revealed a unique spectral signature in OSCC cells at excitation at 360-400 nm light, which is above the hazardous range, and emission at 575-650 nm, with potentially longer Stokes shift. These wavelengths could be leveraged to engineer optical devices that are more precise that currently-used handheld devices for OSCC detection in clinical applications.

Loss of autofluorescence in cancer tissue is clinically observed in patients with OSCC^15, 26^, but understanding the molecular basis is important for optimizing the fluorescent wavelength of screening devices to improve clinical detection and lesion removal. Size-exclusion ultrafiltration indicated that a molecule greater than 10 kDa accounted for differential autofluorescence, excluding small molecules like free FAD or NADH. By integrating spectral and proteomic analyses, we show that the only band showing autofluorescence in the SDS-PAGE gel of cell lysates from OSCC and non-cancer cells was at 70 kDa. This band appears to be responsible for the loss of autofluorescence in multiple OSCC cell lines at the wavelengths that are examined by some commonly-used clinical optical devices (excitation: 400-460 nm; emission: 470-580 nm). High resolution imaging revealed a characteristic punctate pattern, suggesting that the 70 kDa protein is localized to membrane-bound cytoplasmic organelles^17^. Among membrane-bound organelles, flavoproteins are primarily compartmentalized in the mitochondrion^18, 19^, and reports of the limited number of natural fluorophores, altered levels of flavin in cancer cells and similarity in the spectral wavelengths of flavin and flavoproteins^27^ suggest that a flavoprotein is responsible. Of several candidate proteins obtained by LC-MS/MS screening, SDHA and ACADV are mitochondrial flavoproteins with 70 kDa as molecular weight. Transfering the protein from the SDS-PAGE gel to western blot for assessment revealed lowered SDHA levels. More importantly, measuring autofluorescence (within the FAD spectral region) from SDHA (recovered by immunoprecipitation) confirmed reduced fluorescence in OSCC than in non-cancer cells, suggesting flavinated SDHA is responsible for loss of autofluorescence. Unlike unflavinated SDHA^28^, the FAD moiety on flavinated SDHA is responsible for flavinated SDHA being autofluorescent^29^. Previously, more than half of the fluorescent signals from the mitochondrion was shown to be NAD-linked^28^. Therefore, NAD-linked, flavinated α-lipoamine dehydrogenase (approximately 54 kDa) was observed to be the major source of autofluorescence from the rat liver^28^. However, as the band on the SDS-PAGE gel of cell lysates was at 70 kDa, we ruled out flavinated α-lipoamine dehydrogenase (54 kDa) as a candidate.

Flavination of SDHA is intrinsically linked to complex formation with SDHB for function^20^. Consistently, immunohistochemical staining in SDHA-related gastrointestinal tumors reveals normal SDHA but lowered SDHB expression^30, 31^. No loss of SDHA and SDHB staining has also been reported in SDHA-related pheochromocytoma and paraganglioma^32^, and up to 25% of SDHA-related tumors could have normal expression levels^33^. As there are no antibodies specifically for flavinated-SDHA, SDHB levels were used as a surrogate for functional, flavinated, SDHA levels. The comparison of adjacent normal tissues to the tumor tissue in the same tissue sections of multiple patient samples revealed compelling evidence of decreased functional or flavinated SDHA in tumor tissue using SDHB levels as an indicator.

Demonstrating that loss of tissue autofluorescence is related to mitochondrial flavoproteins implies altered cellular metabolism. Supporting this notion, we observed increases in oxidative phosphorylation (OXPHOS) in OSCC which may represent an essential requirement for cancer progression. Previously, functional restoration of OXPHOS in B16 melanoma and 4T1 breast carcinoma has been identified as a prerequisite for efficient tumor formation, invasion and metastasis^34^. Additionally, elevated OXPHOS sustains survival of prostate, colon and breast cancers which are resistant to docetaxel, 5-fluorouracil and aromatase inhibitors, respectively^35–37^. Interestingly, increased OXPHOS in our study correlated with decreased levels of SDHA among OSCC cell lines. SDHA catalyzes the oxidation of succinate to fumarate in the presence of FAD, and lowered expression or function of SDHA will result in accumulation of succinate in the cytosol^38^. Succinate is a potent oncometabolite and inductor of superoxide formation^38, 39^. Therefore, altered function or expression of SDHA has been observed in neuroblastoma, renal carcinoma, pituitary adenoma, paraganglioma-pheochromocytoma syndrome and gastrointestinal stromal _tumors_^30,31,40,41^.

Selectively inhibiting ATP synthesis (synthase) using oligomycin does not abolish differential OXPHOS levels between OSCC and non-cancer cells. In fact, there is still difference in OXPHOS which is only accounted for by inhibiting complex I and III of the electron transport chain (using rotenone and antimycin A). In addition to a role for ATP production, this indicates that oxygen consumption^42, 43^ at complex I and III are critical to the unique, metabolic, mitochondrial signature of OSCC. In support of a role for Complex I in the mitochondrial phenotype of OSCC, we observed a positive relationship between OXPHOS and ACADV intensity in our study, where increased OXPHOS in some cancer cells relative to non-cancer cells was mirrored in ACADV western blot intensity levels. ACADV is required for the assembly and function of Complex I^44^ and our results provide grounds for investigating its metabolic role in OSCC progression. Both complex I and III, but primarily complex I, play important roles in mitochondrial superoxide production from oxygen consumption^45^. Superoxide forms damaging reactive species which, in addition to succinate, could stabilize HIF-1α expression, cause epigenetic changes, drive NF-kB signaling, oxidize amino acid residues, damage DNA and cause genomic instability, and participate in retrograde signaling^38, 46, 47^. Although HIF-1α is a key transcriptional driver of glycolysis, NF-kB signaling and ACADV expression could have maintained OXPHOS^44, 48, 49^, accounting for the concomitant elevation of OXPHOS and glycolysis that we observed in OSCC. Moreover, a positive feedback loop between glycolysis and OXPHOS through O-linked ϕ3-N-acetyl glucosamine modification of IKKB during oncogenesis has been previously noted^50, 51^.

Metabolic heterogeneity characterized by elevated glycolysis in some cancer subpopulations and increased OXPHOS in others has been documented in pancreatic ductal adenocarcinoma (PDAC), breast, prostate, ovarian and peritoneal cancers^52–58^. For example, whereas activation of KRAS drives glycolysis in PDAC^54^, RAGE and HMGB1^55^, HSP 60^56^ and COX6B2^57^ enhance OXPHOS which is required for proliferation, epithelial-mesenchymal transition and metastasis among PDAC subpopulations. However, in our study, OSCC demonstrated functional increases in both glycolysis and OXPHOS among the same cell population. Such metabolic profile is still uncharacterized in cancer biology, and has only been documented in a resistant clone of PDAC cancer stem cells (CSC) where suppression of MYC and increase in PGC-1α underlie increment in both OXPHOS and glycolysis^59^. Importantly, OSCC cells did not reveal a compensatory increase in glycolytic flux following inhibition of mitochondrial respiration by rotenone and antimycin A, suggesting that OSCC may be susceptible to therapies that target OXPHOS. This lack of bioenergetic compensatory mechanisms is an emerging feature of multidrug-resistant melanoma, myeloid leukemia, and PDAC CSC^57, 59–62^. Lastly, proton-linked lactate flux occurs through monocarboxylate transporters (MCTs) in normal and tumor cells^63^, and OSCC cells revealed elevated proton efflux rate (PER). This suggests that MCTs play an important role in OSCC metabolism and autofluorescence, and provides a basis for studies aimed at targeting the MCT pathway in the development of new OSCC therapies.

Conclusively, the intrinsic optical properties of tissues are an indicator of cellular structure, function and metabolism, and can be useful prognostic indicators and predictors of therapeutic outcome. In addition, the unique metabolic changes in OSCC cell lines could be targets for single or combination therapy.

**Supplementary Figure 1.**
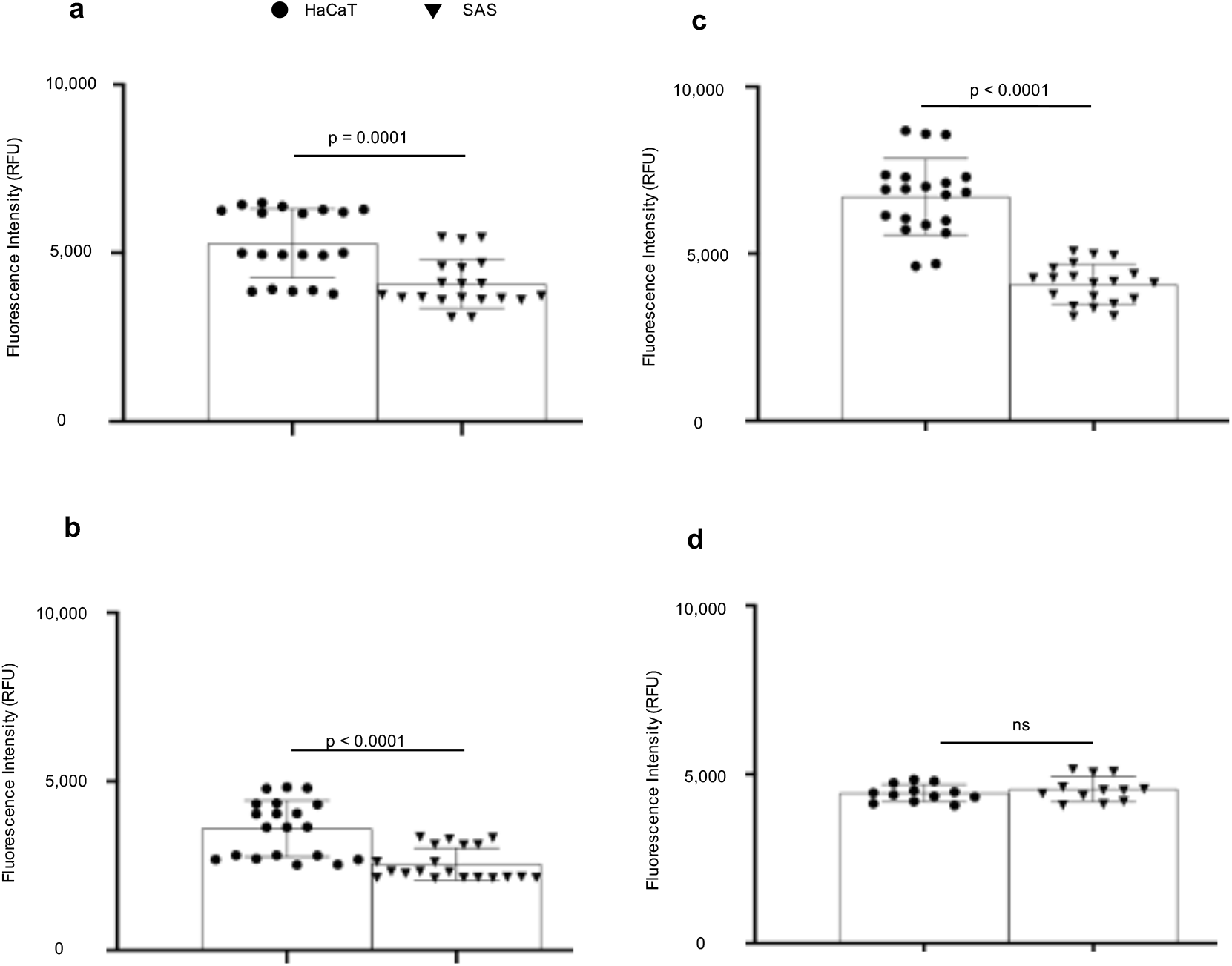
Decreased autofluorescence in oral squamous cell carcinoma (OSCC) cells is likely due to molecules greater than 10 kDa in size. **a-c,** Fluorescence intensities of cell suspension (a), unfiltered supernatant from cell lysates (b) and supernatant filtered through a 30 kDa ultra filter are reduced in SAS cancer cell line compared to non-cancer HaCaT cells. **d,** Fluorescence intensity of supernatant filtered through a 10 kDa Ultra Filter is similar between SAS cancer cell line and non-cancer HaCaT cells. Two-tailed unpaired t-test, n = 20 (a-c), n = 12 (d).

**Supplementary Figure 2.**
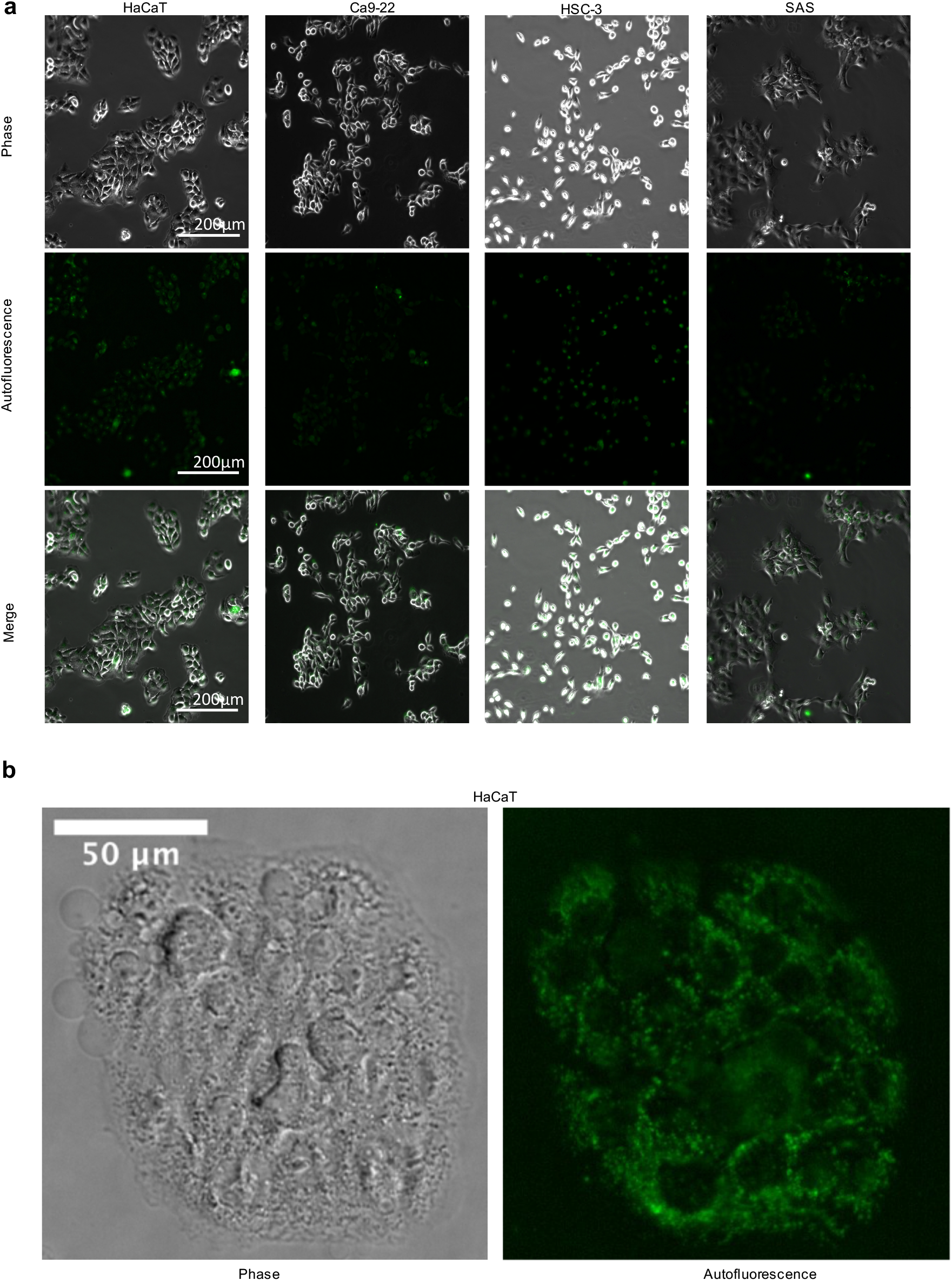
Fluorescence microscopy of unstained cells reveal a punctate pattern in non-cancer (HaCaT) cells. **a,** Imaging by blue light reveals decreases in autofluorescence among oral squamous cell carcinoma (OSCC; Ca9-22, HSC-3 and SAS) when compared to HaCaT cells. **b,** Autofluorescence in HaCaT cells reveal a characteristic punctate pattern.

## Methods

### Cell culture

Human oral squamous cell carcinoma (OSCC) cell lines derived from the gingiva (Ca9-22), tongue (SAS), and a site of lymph node metastasis (HSC-3) were obtained from the Human Science Research Resources Bank (Osaka, Japan). The immortalized normal human epidermal keratinocyte cell line (HaCaT) was obtained from Cell Lines Service (Eppelheim, Germany). All cell lines were cultured at 37° Celsius (C) in 5% CO_2_ using Dulbecco’s Modified Eagle’s Medium (DMEM, Thermo Fisher Scientific, MA, USA) supplemented with 10% fetal bovine serum (FBS) (Biowest, France) and 1% penicillin/streptomycin (Thermo Fisher Scientific, MA, USA).

### Cell lysis

Cells were washed twice with 5 mL of phosphate-buffered saline (PBS). Afterwards, 100 μL of protein RIPA lysis buffer (The Merck Group, Darmstadt, Germany) with proteinase (Halt™ Protease, Thermo Fisher Scientific, MA, USA) and phosphatase inhibitors (Phosphatase Inhibitor Cocktail, 100X, Thermo Fisher Scientific, MA, USA) were added, following the manufacturer’s protocol. Adherent cells were collected by scraping off plastic surfaces in the presence of the lysis buffer. The cell suspension was subjected to two freeze-thaw cycles by placing cells on ice for 30 minutes (min) and at -80℃ for 10-min twice. Lysates were centrifuged at 14800 rpm for 10-min and the supernatant was collected for further analysis.

### Fluorescence spectroscopy

Fluorescence excitation-emission matrices (EEMs) of HaCaT and OSCC (SAS, HSC-3, Ca9-22) cell lysates as well as FAD and NADH (both from Sigma-Aldrich, Missouri, USA, as controls) were measured using a fluorescence spectrophotometer, SpectraMax (R) M3 (Molecular Devices, CA, USA). Samples were diluted with PBS and placed in a non-fluorescent quartz 10 mm cuvette. The protein concentration of each cell lysate was determined, and concentrations were adjusted to 100 μg/ mL for quantitative comparisons, and the concentrations of FAD and NADH were set to 0.1 mg/ mL as previously described^64^. The gain of the photomultiplier tube (PMT) was carefully adjusted to avoid saturation and set to be the same among all measurements. To compensate the background fluorescence and the signal offset of the measurements, the EEM of PBS and RIPA were subtracted from the samples’ EEMs before visualization and analysis^64^. To quantitatively evaluate the fluorescence intensity at wavelengths used by the Illumiscan^®^ (SHOFU, Kyoto, Japan) handheld imaging device (excitation: 400-460 nm; emission: 470-580 nm)^16^, we measured the EEMs from five samples using filters (excitation: 400-450 nm, emission: 490-585 nm). The ranges of excitation and emission were selected to broadly contain the signatures of FAD and NADH as well as any proteins that could bind FAD and NADH and alter the peak absorption and emission spectra of these molecules.

### Size-exclusion ultrafiltration

SAS and HaCaT cells at 1×10^5^cells/ mL of PBS were assessed as cell suspension for fluorescence intensity using a fluorescence spectrophotometer as described above. Similarly, 1×10^5^cells/ mL of PBS were lysed then centrifuged (14,800 x g) for 10 mins to collect unfiltered supernatant, whose fluorescence intensity was evaluated. Subsequently, supernatant was evaluated for fluorescence intensity after being filtered using either a 30 or 10 kDa molecular weight cutoff (MWCO) ultra filter (Vivaspin 500; Cytiva, MA, USA).

### Fluorescence microscopy

HaCaT and OSCC cell lines were cultured on Eppendorf glass chamber slides (Hamburg, Germany) for 3 days (d) and imaged, without any stains, using an inverted fluorescence microscope (Eclipse Ts2R, Nikon, Japan). Prior to imaging, cells were washed 3 times with PBS and replaced with non-fluorescent, FluoroBrite DMEM Medium (Thermo Fisher Scientific, MA, USA). Imaging was performed using Nikon’s CFI Plan Fluor 20x objective (NA0.45) and Lumencor’s SOLA light engine. Fluorescence images were captured with a standard GFP filter cube (excitation: 450-490 nm, emission: 500-550 nm, dichroic mirror).

### Sodium dodecylsulfate polyacrylamide gel electrophoresis (SDS-PAGE) and analysis of autofluorescence

Since we had size-exclusion data indicating that the loss of autofluorescence was not due to unbound small molecules (e.g. FAD), we looked at the contribution of proteins, specifically flavoproteins. To determine the molecular weight of autofluorescent molecule(s) in the cell lyses, each lysate containing 50 μg of protein was examined using a modified SDS-PAGE, relative to a protein ladder (Prestained Protein Standards, Bio-Rad, Hercules, CA, USA). Proteins in the lysates were suspended in loading buffer containing SDS but were not heated, and then the proteins were separated by PAGE. After the separation, the unstained gels were imaged with filtered UV light using specific wavelengths of illumination (Epi-blue, excitation: 460-490 nm, emission: 518-546 nm) on a gel imager (ChemiDoc MP Imaging System, Bio-Rad, Hercules, CA, USA).

### Western blot analysis

Western blotting was performed to confirm the expression of candidate proteins in the cell lysates. After the SDS-PAGE, the separated proteins were transferred to Mini-size LF PVDF membrane (Bio-Rad Laboratories, Hercules, CA, USA). The membranes were blocked for 1-hour (h) using condensed milk 1%. Thereafter, the membranes were incubated overnight at 4°C with an anti-ACADV polyclonal antibody (Novus Biologicals, Centennial, CO, USA) and an anti-SDHA monoclonal antibody (Cell Signaling Technology, Danvers, MA, USA), along with anti-GAPDH (Glyceraldehyde 3-phosphate dehydrogenase) antibody (Proteintech Group Inc, Rosemont, IL, USA) as an internal control. The membranes were then incubated for 2-h at room temperature with rabbit HRP-linked IgG (GE Healthcare, Chicago, IL, USA), and a chemiluminescence detection reagent (SuperSignal™ West Pico PLUS Chemiluminescent Substrate, Thermo Fisher Scientific, MA, USA) was added to detect binding of the secondary antibody. Imaging and analysis were performed with the ChemiDoc MP Imaging System.

### Liquid chromatography with tandem mass spectrometry (LC/MS/MS)

LC/MS/MS analyses were performed at Michigan State University mass spectrometry facility. Bands containing candidate proteins with an autofluorescent signature were cut out of SDS-PAGE gels and digested in-gel according to a published protocol with modifications^65^. For scientific rigor, a batch of HaCaT lysate (standard) was compared to the HaCaT (and OSCC cell) lysates used during LC-MS/MS. Peptides were extracted from the gel and vacuum dried to 2μl were then re-suspended in 100μl of 100mM triethyl ammonium bicarbonate (TEAB) and labeled with Tandem Mass Tag (TMT) reagents (Thermo Fisher Scientific, MA, USA) according to manufacturers’ instructions. The remaining samples were combined in equal amounts, by volume, and this mixture was purified by solid phase extraction. Eluted peptides were dried by vacuum centrifugation to ∼2μl and stored at -20°C. Dried samples were re-suspended in 20μl of 2% acetonitrile/0.1% trifluoroacetic acid. An injection of 10-μl was automatically made using EASYnLC 1000 (Thermo Fisher Scientific, MA, USA) onto HPLC Columns (Acclaim PepMap RSLC 0.075mm x 20mm C18 trapping column, Thermo Fisher Scientific, MA, USA) and washed for ∼5-min with buffer A (99.9% Water/0.1% Formic Acid). Bound peptides were then eluted over 125-min with a gradient of 8% Buffer B (99.9% Acetonitrile/0.1% Formic Acid) to 40% Buffer B in 114-min, ramping to 90% Buffer B at 115-min and held at 90% Buffer B for the duration of the run at a constant flow rate of 300-nl/min. Column temperature was maintained at a constant temperature of 50°C using and integrated column oven (PRSO-V1, Sonation GmbH, Biberach, Germany). Eluted peptides were sprayed into Q-Exactive mass spectrometer (Thermo Fisher Scientific, MA, USA) using a FlexSpray spray ion source. Survey scans were taken in the Orbi trap (140000 resolution, determined at m/z 200) and the top twelve ions in each survey scan are then subjected to automatic higher energy collision induced dissociation (HCD) with fragment spectra acquired at 70000 resolution. The resulting MS/MS spectra are converted to peak lists using Proteome Discoverer, v2.2 (Thermo Fisher Scientific, MA, USA) and searched against all human protein sequences available from Uniprot (downloaded from www.uniprot.org on 2017-11-02) appended with common laboratory contaminants (downloaded from www.thegpm.org, cRAP project) using both the Sequest and Mascot searching algorithms. The output was then analyzed using Scaffold, v4.8.9 (www.proteomesoftware.com) to probabilistically validate protein identifications. Assignments validated using the Scaffold 1% FDR confidence filter are considered true.

### Immunocytochemistry (ICC)

The following staining was done according to the protocol specified by Cell Signaling Technology. Briefly, HaCaT and OSCC cells were cultured on glass chamber slides (Eppendorf, Hamburg, Germany) for 3 d and stained with anti-SDHA monoclonal antibody (Cell Signaling Technology, MA, USA) and anti-ACADV polyclonal antibody (Novus Biologicals, CO, USA) and imaged with an inverted fluorescence microscope (Nikon, Japan). The culture medium was removed and the cells were washed 3 times with PBS. Cell samples on slides were blocked with blocking buffer for 60 min at room temperature. The blocking buffer was removed and each diluted primary antibody (anti-ACADV: 2 μg/mL, anti-SDHA: 10 μg/mL) was applied to samples and incubated overnight at 4°C. After incubating, samples were rinsed and incubated for 1-h with secondary antibody before imaging in the Nikon fluorescence microscope.

### Subjects’ demographics, SDHA and SDHB immunohistochemistry (IHC) and quantification

Sequential tissue slides were obtained from research tissues collected from 5 subjects enrolled under NCT02415881. This study protocol was approved by the Stanford University Institutional Review Board (IRB-35064) and the FDA (NCT02415881) with written informed consent obtained from all patients. The study was performed in accordance with the Helsinki Declaration of 1975 and its amendments, FDA’s ICH-GCP guidelines, and the laws and regulations of the United States. Briefly, subjects’ age ranged 32-78 years and 20% was female. All subjects were diagnosed with primary head and neck squamous cell carcinoma located on the lateral tongue (n = 2), the floor of the mouth (n = 1), in the larynx (n = 1) or on the alveolar ridge (n = 1). Subject’s tumor stage was cT3 in 2 cases, cT4a in 2 cases, and one patient presented with recurrent larynx cancer that was previously treated with chemo-radiation (cT-stage unknown). Pathological cancer stage was pT3 in 1 subject, pT4a in two subjects and pT2 in 2 subjects. None of the subjects reported on alcohol abuse, and 2 out of 5 subjects never smoked, one subject quit smoking and two subjects reported to be current smokers. Three subjects were HPV negative, one positive and one unknown. To examine the expression of SDHA and SDHB, automated immunohistochemistry (IHC) staining was performed on sequential sections using a Dako Autostainer (Agilent Technologies, Santa Clara, CA, USA) for SDHA (1:250; D6J9M; Cell Signaling), Danvers, MA), and SDHB (1:250; Ab14714; Abcam, Cambridge, MA). For SDHA, no secondary antibody was used. With SDHB, Envision FLEX+ rabbit (linker) (prediluted, SM805, Agilent Technologies) was used. Positive and negative controls were included in each staining batch. Immunoreactivity was visualized with diaminobenzidine and magenta chromogens (Dako EnVision, Glostrup, Denmark). Digital images of IHC-stained slides were obtained at 4 to 20x magnification with a whole slide scanner (NanoZoomer 2.0-HT slide scanner; Hamamatsu Photonics, Hamamatsu City, Japan). For comparison, H&E slides were obtained on which tumor was outlined by a pathologist. For quantification of SDHB expression, regions of interest were randomly chosen. In 4 of 5 patients, tumor-negative and tumor-positive areas were present on the evaluated slides. In the remainder patient (patient 4), no tumor-positive tissue was present; therefore, this subject was excluded from further analysis. Combined, a total of 6 paired areas of tumor-negative and tumor-positive tumor areas were evaluated as follows: 2 pairs each in patient 5 and patient 3, 1 pair each in patient 1 and patient 2. Pathologist-based tumor regions were annotated on slides using Aperio’s annotation software (ImageScope Viewing Software: Positive Pixel Count v9.1, Aperio ImageScope®; Leica Microsystems Inc.) and the intensity of staining was graded as follows: negative, weak positive (Intensity Threshold weak [upper limit] = 220, [lower limit] = 175), medium ([upper] = 175, [lower] = 100), and strong ([upper] = 100, [lower] = 0) by default. The staining of SDHB was then quantified by IHC positivity, which was calculated as the number of positive pixels stained at each positive intensity level divided by the total number of pixels (the number of positive and negative pixels).

### Subject fluorescence imaging

As an example, the Illumiscan was used to examine loss of fluorescence intensity in an 86-year old female diagnosed with T1 N0 M0 tongue cancer on the right side (Fig. 1). The study protocol was approved by the ethics committees of Tokyo Dental College (Approval number: 740), after obtaining patient consent. The study was performed in accordance with the Helsinki Declaration of 1975 and its amendments.

### Immunoprecipitation (IP)

IP was performed according to the manufacturer’s (Cell Signaling Technology, MA, USA) protocol. Summarily, 4 μL of primary anti-SDHA antibody was added to 200 μg/mL cell lysate and incubated overnight at 4 °C. Previously washed magnetic beads (Cell Signaling Technology, MA, USA) were mixed with cell lysate and incubated at room temperature for 20-min. The beads in the solution were separated using a magnetic separation rack, and the pellet was washed five times on ice with RIPA buffer, followed by protein elution at 95 °C. Purified SDHA was detected in the Molecular Devices SpectraMax M3 spectrophotometer.

### Live-cell metabolic assay

Real-Time ATP rate assay of oral carcinoma cell lines (SAS, HSC-3 and Ca9-22) in comparison to normal keratinocytes (HaCaT) was determined using Seahorse XFp as previously described^66, 67^. For each cell line, 50,000 cells were seeded in complete DMEM containing 1% penicillin-streptomycin for 12 h. One hour prior to running the assay, full media was replaced by unbuffered XF assay media (pH 7.4) supplemented with 25 mM D-glucose and 4 mM L-glutamine, and kept in a non-CO_2_ incubator at 37°C. Final well concentrations of 1.5μM and 0.5μM of oligomycin and rotenone/antimycin A, respectively, were sequentially injected after obtaining baseline measurements. Oxygen consumption rates (OCR), extracellular acidification rate (ECAR), proton efflux rate (PER) were repeatedly measured, and expressed as mean (SD). The materials and drugs used for the assay were sourced from Seahorse Bioscience (MA, USA).

### Statistics and reproducibility

Data were expressed and analyzed as indicated in respective figure legends using using GraphPad Prism version 8.4.3 (GraphPad Software, San Diego CA; www.graphpad.com). For comparative analysis of proteins extracted from gels by LC/MS/MS, Scaffold, version 4.8.9 (www.proteomesoftware.com) was used. Statistical significance was set at p < 0.05.

### Data availability

The data supporting the findings of this study are available within the paper and its Supplementary Information.

## Acknowledgements

Funding for this work was provided in part by the James and Kathleen Cornelius Endowment at MSU; and a grant from the NIH/NCI to C.H.C. (R01 CA182043). Dr. Takeshi Onda, Dr. Tadashi Miura, and Dr. Koji Tanabe of Tokyo Dental College assisted with mass spectrometry and provided expertise that guided the design of some experiments. Max M. Kuhnert of Michigan State University made the Supplementary Tables.

## Author contributions

Conceptualization, T.M., C.V.M., T.S. and C.H.C.; Methodology, T.M., C.V.M., T.S. and C.H.C.; Investigation, T.M., C.V.M., E.U., E.H. A., S. J. C., N.S.V., Q.Z., B.A.M., E.L.R., T.S. and C.H.C.; Writing – Original Draft, T.M. and C.V.M.; Writing – Review & Editing, T.M., C.V.M., E.U., E.H. A., S. J. C., N.S.V., Q.Z., B.A.M., E.L.R., T.S. and C.H.C.; Funding Acquisition, C.H.C.; Resources, E.L.R. and C.H.C.; Supervision, T.S. and C.H.C.

