## Supplementary Table 1 for "Flavinated SDHA Underlies the Change in Intrinsic Optical Properties of Oral Cancers"

**Supplementary Table 1**: Sixteen proteins of approximately 70kDa were identified using LC-MS/MS from the distinct autofluorescent band of HaCaT (non-cancer) cell lysates, and these proteins were reduced (%) in oral squamous cell carcinoma (Ca9-22, HSC-3, SAS) cell lysates.

| No | Name | Accession Number | Alternate ID | Molecular weight (kDa) | HaCaT | Ca9-22 | HSC-3 | SAS | Localized |
|---|---|---|---|---|---|---|---|---|---|
| 1 | Keratin, type II cytoskeletal 1 OS=Homo sapiens OX=9606 GN =KRT1 PE=1 SV=6 | K2C1_HUMAN(+4) | KRT1 | 66 | 3% (0.0) | -20%(-0.2) | -4%(0.0) | -98%(-1.0) | Not available |
| 2 | Phenylalanine—tRNA ligase beta subunit OS=Homo sapiens OX=9606 GN=FARSB PE=1 SV=1 | SYFB_HUMAN(+2) | FARS8 | 66 | 1%(0.0) | -202% (-2.0) | -102%(-1.0) | -99%(-1.0) | Nucleoplasm |
| 3 | Apoptosis-inducing factor 1, mitochondrial OS=Homo sapiens OX=9606 GN=AIFM1 PE=1 SV=1 | AIFM1_HUMAN | AIFM1 | 67 | 0%(0.0) | -3%(0.0) | -41%(-0.4) | -36%(-0.4) | Not Available |
| 4 | Calnexin OS=Homo sapiens OX=9606 GN=CANX PE=1 SV=2 | CALX_HUMAN | CANX | 68 | 1%(0.0) | -108%(-1.1) | -25%(-0.2) | -22%(-0.2) | Endoplasmic reticulum |
| 5 | V-type proton ATPase catalytic subunit A OS=Homo sapiens OX=9606 GN=ATP6V1A PE=1 SV=2 | VATA_HUMAN(+1) | ATP6VIA | 68 | 0%(0.0) | -146%(-1.5) | -25%(-0.3) | -45%(-0.4) | Nucleoplasm |
| 6 | Dolichyl-diphosphooligosaccharide—protein glycosyltransferase subunit 2 OS=Homo sapiens OX=9606 GN=RPN2 PE=1 SV=3 | RPN2_HUMAN (+1) | RPN2 | 69 | -1%(0.0) | -34%(-0.3) | -46%(-0.5) | -41%(-0.4) | Nucleoplasm |
| 7 | Xaa-Pro aminopeptidase 1 OS=Homo sapiens OX=9606 GN=XPNPEP1 PE=1 SV=3 | XPP1_HUMAN | XPNPEP1 | 70 | 0%(0.0) | -6%(-0.1) | -7%(-0.1) | -18%(-0.2) | Cytosol |
| 8 | Very long-chain specific acyl-CoA dehydrogenase, mitochondrial OS=9606 GN=ACADV PE=1 SV=1 | ACADV_HUMAN(+2) | ACADV | 70 | 1%(0.0) | -155%(-1.5) | -88%(-0.9) | -46%(-0.5) | Nucleoli and mitochondria |
| 9 | Cluster of Heterogenous nuclear ribonucleoprotein Q OS=Homo sapiens OX=9606 GN=SYNCRIP PE=1 SV=2 (HNRPQ_HUMAN) | HNRPQ_HUMAN[2] | SYNCRIP | 70 | -0%(0.0) | -57%(-0.6) | -22%(-0.2) | -5%(0.0) | Nucleoplasm |
| 10 | Eukaryotic translation initiation factor 3 subunit L OS=Homo sapiens OX=9606 GN=EIF3L PE=1 SV=1 | BOQY89_HUMAN(+2) | EF3L | 71 | 0%(0.0) | -100%(-1.0) | -85%(-0.8) | -24%(-0.2) | Nucleoli |
| 11 | Succinate dehydrogenase [ubiquinone] flavoprotein subunit, mitochondrial OS=Homo sapiens OX=9606 GN=SDHA PE=1 SV=2 | SDHA_HUMAN | SDHA | 73 | -1%(0.0) | -317%(-3.2) | -162%(-1.6) | -134%(-1.3) | Mitochondria |
| 12 | ATP-dependent DNA Helicase Q1 OS=Homo sapiens OX=9606 GN=REQCL PE=1 SV=3 | RECQ1_HUMAN | RECCL | 73 | -2%(0.0) | -423%(-4.2) | -79%(-0.8) | -193%(-1.9) | Nucleoplasm |
| 13 | Cluster of Aminopeptidase B OS-Homo sapiens OX=9606 GN=RNPEP PE=1 SV=2 (AMPB_HUMAN) | AMPB_HUMAN[2] | RNPEP | 73 | -1%(0.0) | -95%(-1.0) | -33%(-0.3) | -40%(-0.4) | Golgi Apparatus |
| 14 | Cluster of Prelamin-A/C OS=Homo sapiens OX=9606 GN=LMNA PE=1 SV=1 (LMNA_HUMAN) | LMNA_HUMAN[2] | LMNA | 74 | -2%(0.0) | -108%(-1.1) | -81%(-0.8) | -129%(-1.3) | Nucleoplasm |
| 15 | Aspartate—tRNA ligase, mitochondrial OS=Homo sapiens OX=9606 GN=DARS2 PE=1 SV=2 | SYDM_HUMAN(+1) | DAR52 | 74 | 0%(0.0) | -46%(-0.5) | -41%(-0.4) | -21%(-0.2) | Mitochondria |
| 16 | Carnitine O-palmitoyltransferase 2, mitochondrial OS=Homo sapiens OX=9606 GN=CPT2 PE=1 SV=2 | CPT2_HUMAN | CPT2 | 74 | 2%(0.0) | -104%(-1.0) | -46%(-0.5) | -40%(-0.4) | Nucleoplasm, Nucleoli, Mitochondria |
