## Supplementary Table 1 for "Flavinated SDHA Underlies the Change in Intrinsic Optical Properties of Oral Cancers"

**Supplementary Table 2:** SDHA and ACADV were the only membrane-bound 70kDa flavoproteins present in the autofluorescent band of HaCaT (non-cancer) cell lysates, and were reduced in oral squamous cell carcinoma (Ca9-22, HSC-3, SAS) cell lysates by 46-317%.

| No. | Name | Accession Number | Alternate ID | Molecular weight (kDa) | HaCat | Ca9-22 | HSC-3 | SAS | Localized |
|---|---|---|---|---|---|---|---|---|---|
| 1 | Very long-chain specific acyl-CoA dehydrogenase | ACADV_HUMAN | ACADV | 70 | 1% | -155% | -88% | -46% | Mitochondria |
| 2 | Succinate dehydrogenase [ubiquinone] flavoprotein subunit | SDHA_HUMAN | SDHA | 73 | 1% | -317% | -162% | -134% | Mitochondria |
